# Immobilized enzyme-assisted production of recombinant P113 peptide

**DOI:** 10.64898/2026.04.11.717888

**Authors:** Josephine A. Kirkendoll, Leticia Targino Campos, Emma G. Taylor, Ramiro M. Murata, Robert M. Hughes

## Abstract

Recombinant peptide production was pioneered in the 1970’s for the generation of therapeutic peptides, with notable examples including insulin and somatostatin. These early methods required the use of cyanogen bromide (BrCN) for cleavage of the native peptide sequence from a fusion protein. Since that time, while numerous BrCN-dependent peptide methods continue to be reported, the accessibility and cost of site-specific proteases have improved dramatically. These developments have enabled alternative approaches to recombinant peptide generation that obviate the need for BrCN, an environmentally destructive toxin. We recently created an immobilized SUMO protease that can replace BrCN usage in recombinant peptide production workflows by releasing native peptides expressed as part of a SUMO-peptide fusion protein. We have used this approach to generate P113 peptide, the minimal active fragment of the antifungal peptide Histatin 5. In this report, we describe the creation and characterization of this immobilized SUMO protease and its application in the production of experimentally viable quantities of active P113 peptide.

## INTRODUCTION

Recombinant peptide production has a long history in therapeutic peptide production. Early landmarks in recombinant peptide syntheses include the production of active somatostatin (1977) and insulin (1979) peptides (Itakura et al., 1977; Riggs & Itakura, 1979). These early syntheses were accomplished in an era without readily accessible methods for peptide secretion from host organisms or cell lines and without many of the engineered site-specific proteases that are readily abundant today (1). As such, these strategies used fusion proteins as carriers for peptide sequences and relied upon cyanogen bromide (BrCN) for cleavage of the peptide from the fusion protein (1–3). While effective, this strategy has its limitations, including the required presence of a single internal methionine residue for BrCN cleavage (thus limiting which peptide sequences can be produced by this method) and the toxicity and environmental hazards associated with BrCN usage (4). Despite the ready availability of alternate methods such as solid phase peptide synthesis, BrCN-assisted methods are still in use today (5–7). Reasons for this continued practice include the recombinant production of isotopically labeled peptides (8), the need for post-translationally modified peptides (9), and the use of amber-codon suppression strategies to introduce non-natural amino acids in recombinant systems (10). The goal of this work is to update the traditional BrCN-dependent recombinant peptide workflow with an immobilized enzyme-based strategy (11–13). This method facilitates recombinant peptide production without the use of BrCN while still enabling production of the native peptide sequence. We propose this method as an alternative to BrCN use and as a greener approach to peptide production than solid phase peptide synthesis. The peptide sequence produced in this study (P113) is the minimal 12-amino acid fragment of Histatin 5, a member of the anti-microbial Histatin peptide family (14–17). Naturally present in human saliva, these peptides are a potential source of novel peptide therapeutics that could be applied in the treatment and prevention of gingivitis, dental caries, and Candida infections (14, 17). In particular, they may represent a way to treat drug resistant oral Candida infections in HIV patients, which remain a persistent public health concern (18). The development of new methods for Histatin peptide production has the potential to reignite interest in this demanding area of therapeutic need.

## RESULTS AND DISCUSSION

Traditional BrCN-based methods of recombinant peptide production have followed a consistent series of steps: (1) expression of a fusion protein containing a leader protein or peptide (typically in E. coli) followed by a Met residue and the sequence of the peptide of interest; (2) partial purification of the fusion protein, dialysis, and subsequent cleavage with BrCN; (3) lyophilization of the BrCN cleavage solution, reconstitution, and subsequent purification of the peptide fragment by HPLC; and (4) peptide characterization by mass spec and bioassay (5, 8, 19, 20). In this work, we emphasize that BrCN can be replaced with site-specific proteolysis while still leaving the native peptide sequence intact.

Furthermore, by using an immobilized protease that can be recovered and reused as needed, the efficiency of the process is improved (21). In our method (**Fig. 1**), the peptide of interest is expressed as a fusion to a SUMO tag; this tag is cleaved by SUMO protease after its Gly-Gly C-terminus (22). Barring the inclusion of any additional residues, this cleavage leaves the native peptide sequence intact. To facilitate this process, we created an immobilized SUMO protease using the SENP1 protease fused to HaloTag (23) (**Fig. 2**). Following protocols previously developed in our group (24, 25), we expressed, purified, and immobilized this sequence on magnetic beads and investigated its properties using a commercially available multi-protease substrate (SUMO3-GFP; **Fig. 3**). This substrate enables the comparison of multiple protease activities, which is useful for comparing protease specificity. We compared our previously characterized immobilized TEV protease with the new SUMO protease (**Fig. 3A-B**), confirming that the proteases gave the expected cleavage patterns for each cleavage site. Furthermore, we confirmed that the immobilized protease preparations did not introduce additional material detectable by SDS-PAGE into the assay buffer (**Fig 3B**, rightmost lanes). Finally, to confirm that our SUMO protease preparation was active over multiple trials, we subject an aliquot of protease beads to 4 consecutive cleavage reactions (**Fig. 3C**), confirming that the beads did not lose activity after multiple uses.

**FIGURE 1.**
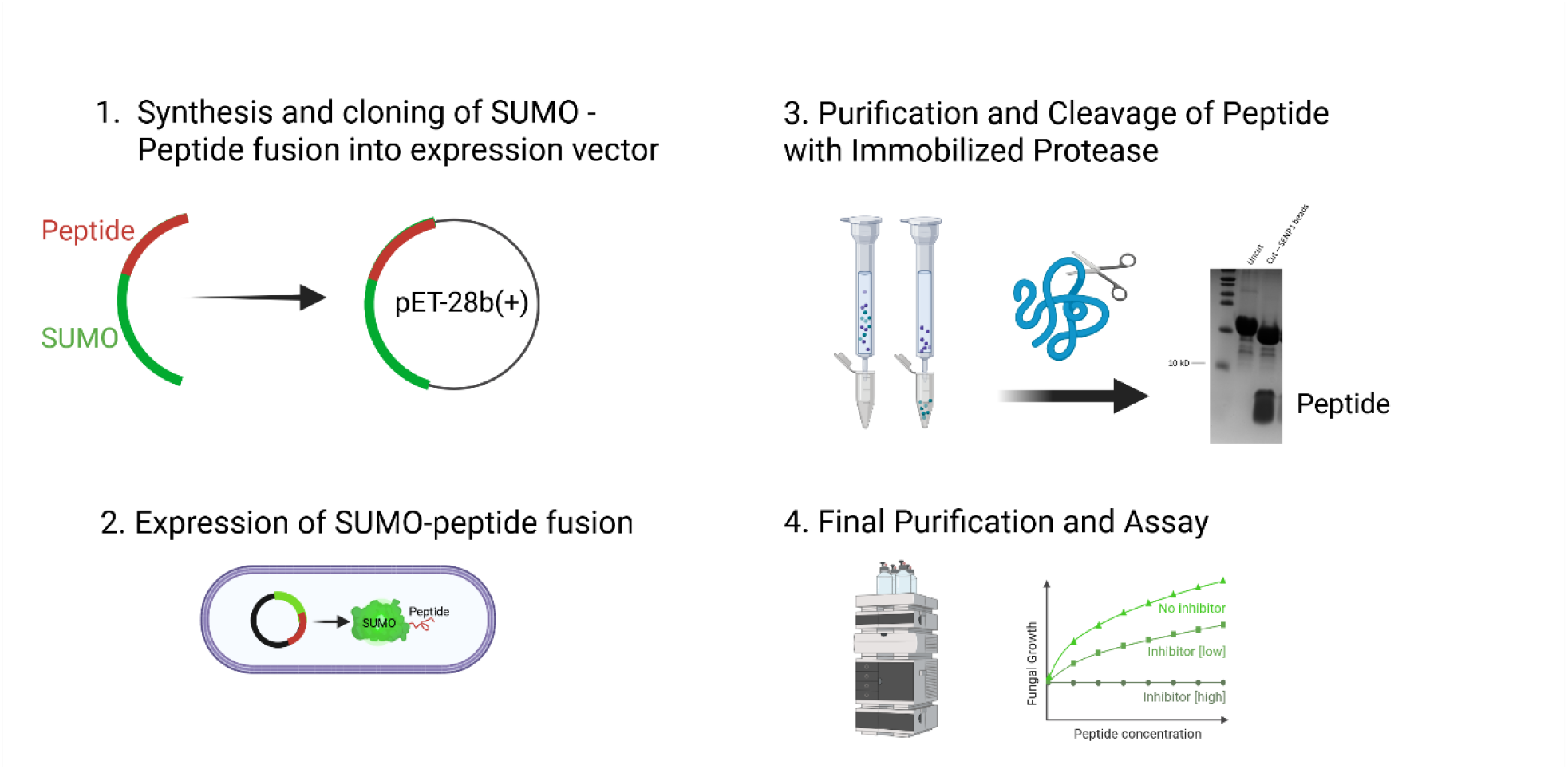
Recombinant peptide production workflow. 1. A gene encoding a SUMO-peptide fusion construct is assembled and cloned into a T7-promoter driven expression vector. 2. SUMO-peptide fusion is expressed in *E. coli*. 3. Purified SUMO-peptide fusion construct is cut with an immobilized SUMO protease, rendering the native peptide sequence. The immobilized protease can be recovered after this step for reuse. 4. Final purification of the recombinant peptide by RPHPLC is followed by mass spectrometry and confirmation of bioactivity.

**FIGURE 2.**
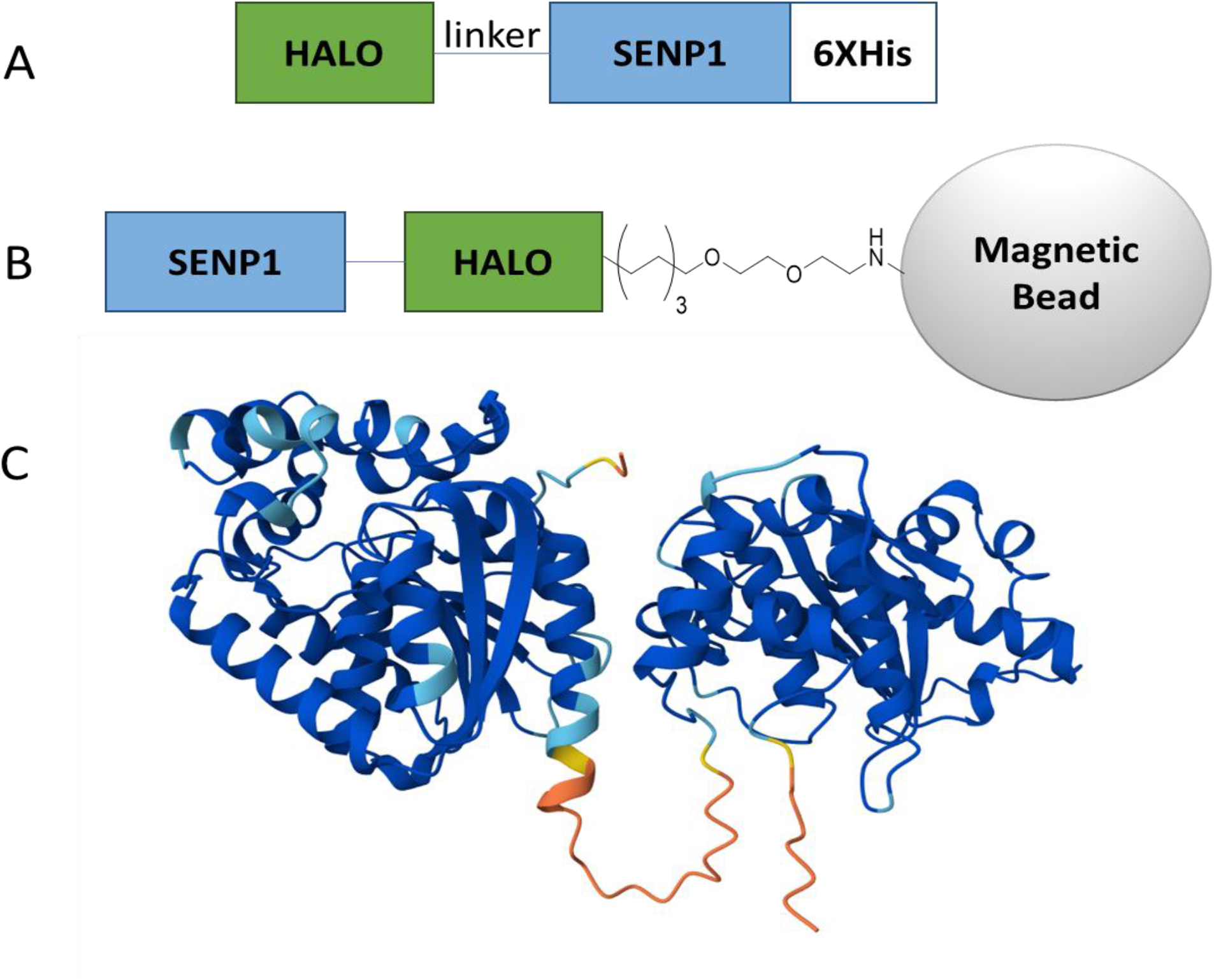
Design and computational model of SUMO protease fusion. A. Schematic of the SUMO protease construct used in this study. HALO = HaloTag protein; linker = GGSD repeat sequence; SENP1 = C-terminal domain of human sentrin/SUMO-specific protease 1; 6XHis = polyhistidine tag. B. Schematic of covalent immobilization of HALO on Halo ligand functionalized magnetic beads. C. AlphaFold3 model of the HaloTag-linker-SUMO-His6X protein. Protein domains are color coded by likelihood of having the predicted structure (pLDDT or predicted local distance difference test) with blue indicating high confidence. Domains appear in same order as in schematic 2A.

**FIGURE 3.**
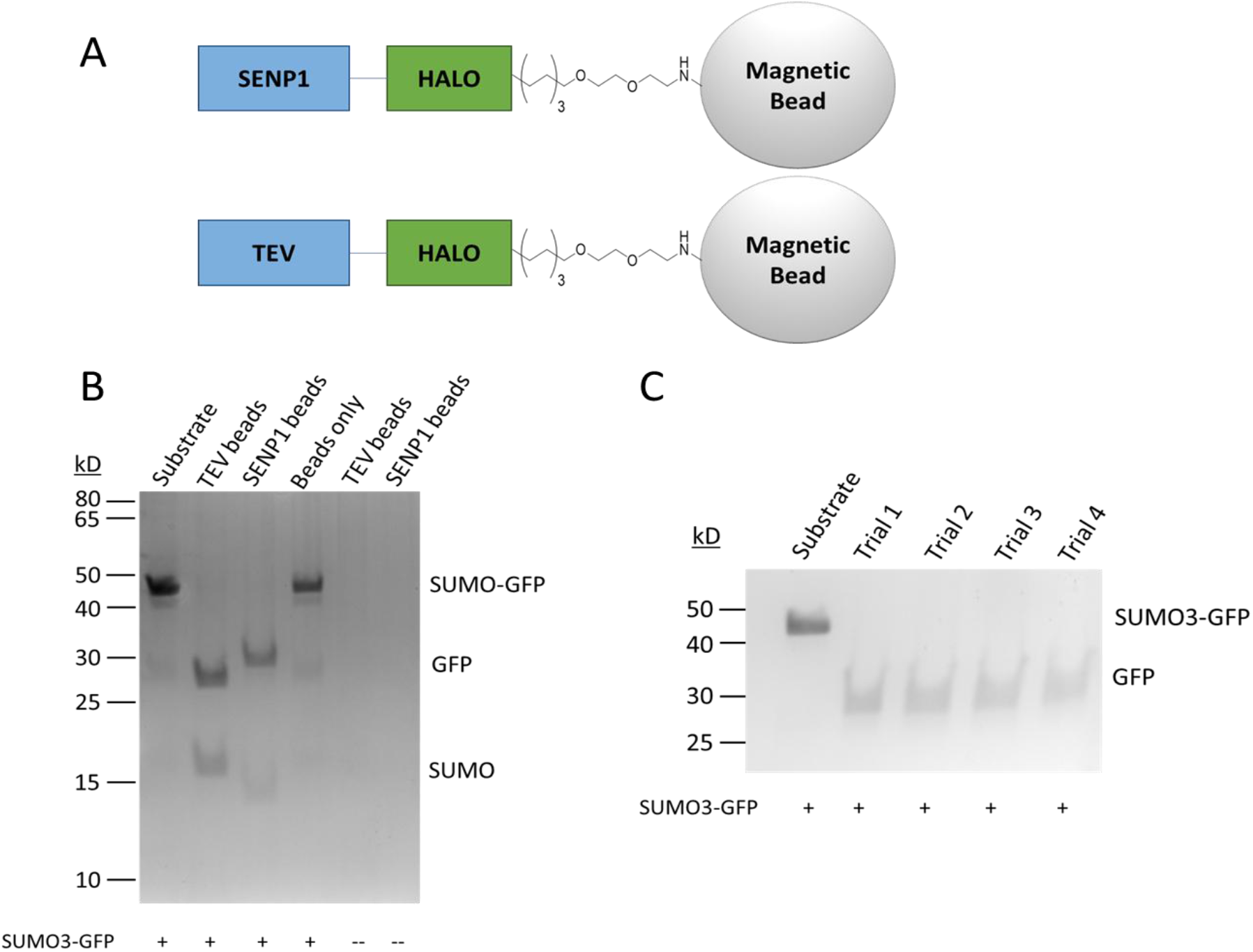
Validation of Immobilized SUMO protease. A. Schematic of HaloTag - immobilized protease constructs. B. SDS-PAGE gel of SUMO-GFP multi-protease substrate with or without incubation of immobilized proteases. The cleavage patterns resulting from TEV and SUMO proteases are consistent with highly specific protease activities. C. SDS-PAGE gel demonstrating repeat use of a single aliquot of immobilized SUMO protease magnetic beads. Consistent high efficiency cleavage of substrate is evident throughout each successive trial.

Having confirmed that the HaloTag-SENP1 labeled beads retained the expected activity profile, we then tested our beads in a recombinant peptide production system. To enable production of the native P113 sequence, a His6X-SUMO-P113 fusion protein was designed (**Fig. 4**). This construct was expressed and purified in E. coli and subsequently tested for its ability to be cleaved by the immobilized SUMO protease beads (**Fig. 5**). These tests demonstrated that the immobilized protease produced a cleavage product (resolved on an 18% SDS-PAGE gel) identical to that produced with non-immobilized protease (**Fig. 5B**), and a time course experiment revealed that while the substrate was mostly cleaved after a 4h incubation period at room temperature, maximum cleavage was achieved after overnight incubation (**Fig. 5C-D**). We also confirmed the robustness of the immobilized protease cleavage by subjecting an aliquot of beads to 10 successive overnight cleavage trials (**Fig. 6**), demonstrating complete proteolysis in each trial. We note that SUMO protease may be an ideal immobilized protease (26, 27) due to the relatively small size of its catalytic domain, its tendency to express in E. coli at high levels in soluble form, and its activity in a wide range of pH and buffer types (28). In our results, the inclusion of a HaloTag to this system does not appear to have compromised the robust nature of the protease, perhaps adding tolerance to protein fusions as another ideal feature of this protease.

**FIGURE 4.**
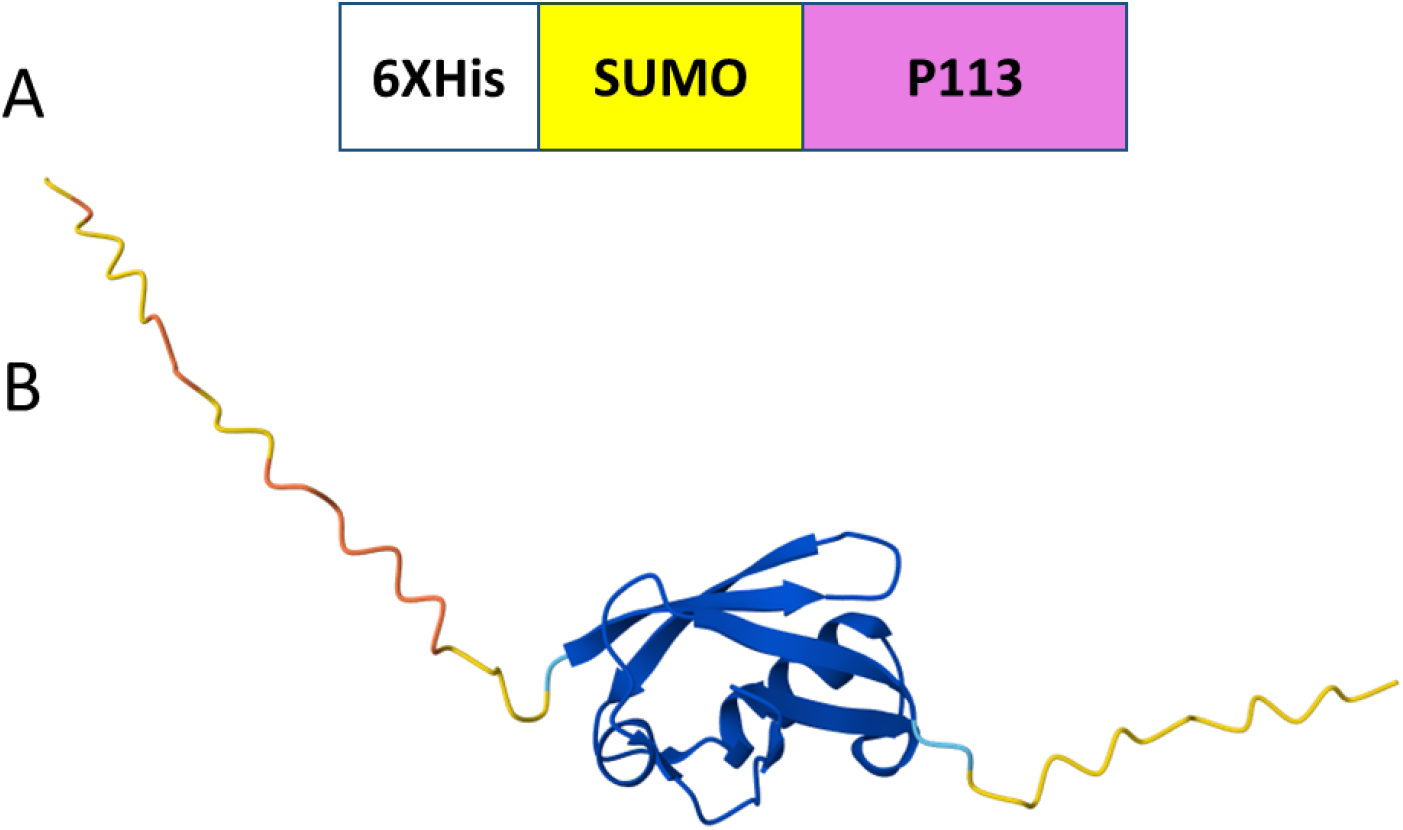
Design and computational model of SUMO – P113 fusion. A. Schematic of the SUMO tag – peptide fusion used in this study. 6XHis = polyhistidine tag; SUMO = Sumo tag; P113 = 12-amino acid fragment of Histatin 5 peptide. B. AlphaFold3 model of the His6X – SUMO tag – P113 protein fusion. Protein domains are color coded by likelihood of having the predicted structure (pLDDT or predicted local distance difference test) with blue indicating high confidence. Domains appear in same order as in schematic 4A. A portion of the N-terminus of the SUMO tag is predicted to be unfolded, hence the longer unstructured region on the left side of the model.

**FIGURE 5.**
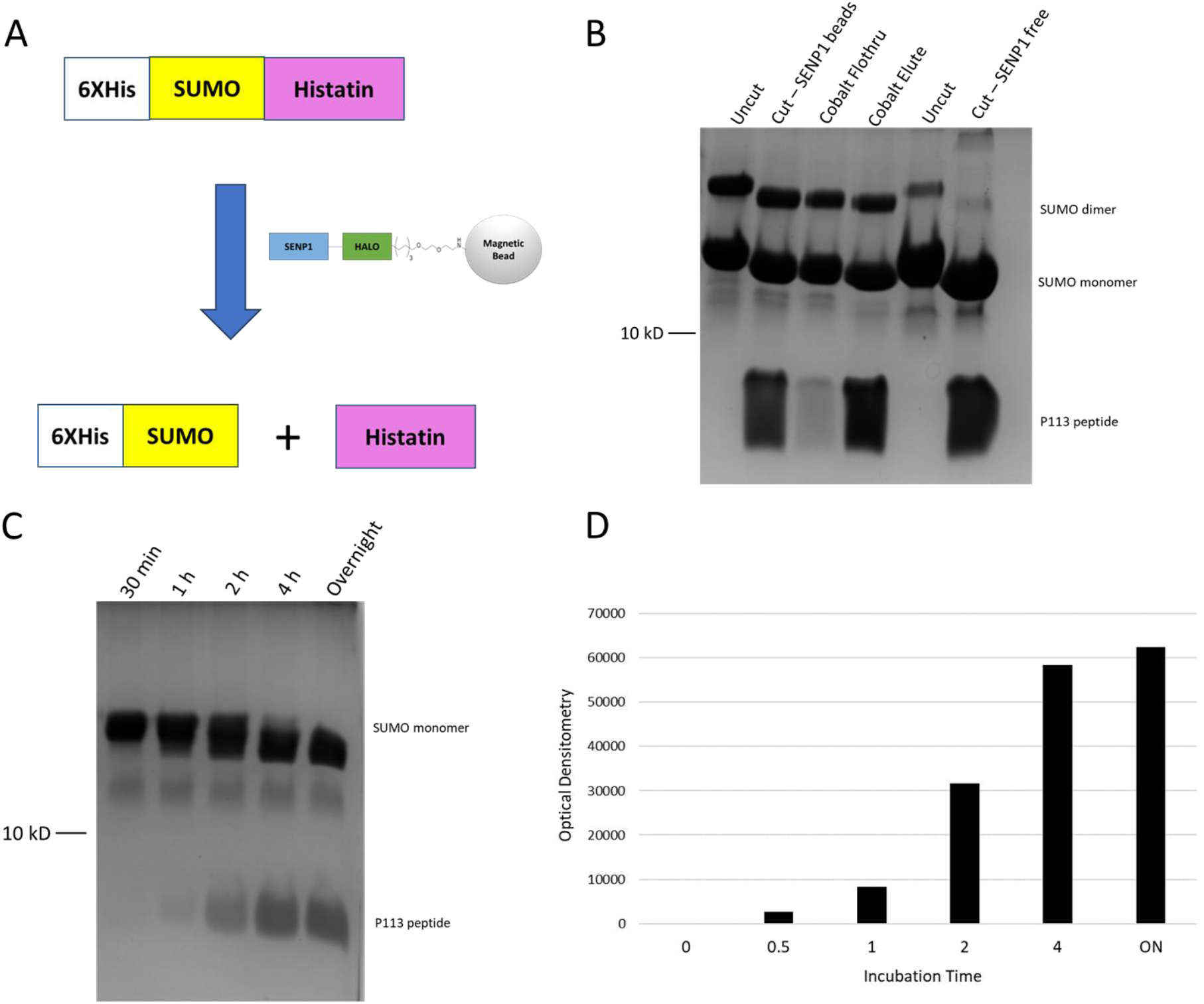
Cleavage of SUMO-P113 peptide with free and immobilized enzyme. A. Schematic of SUMO protease cleavage reaction. B. 18% SDS-PAGE gel showing production of peptide fragments in presence of immobilized SENP1 beads and in the presence of non-immobilized SENP1. The histatin peptide has a high affinity for cobalt resin and is retained alongside the 6XHis tagged SUMO fragment. C. Time course of incubation of SUMO-P113 substrate with SUMO protease beads. D. Optical densitometry of P113 peptide bands in 5C.

**FIGURE 6.**
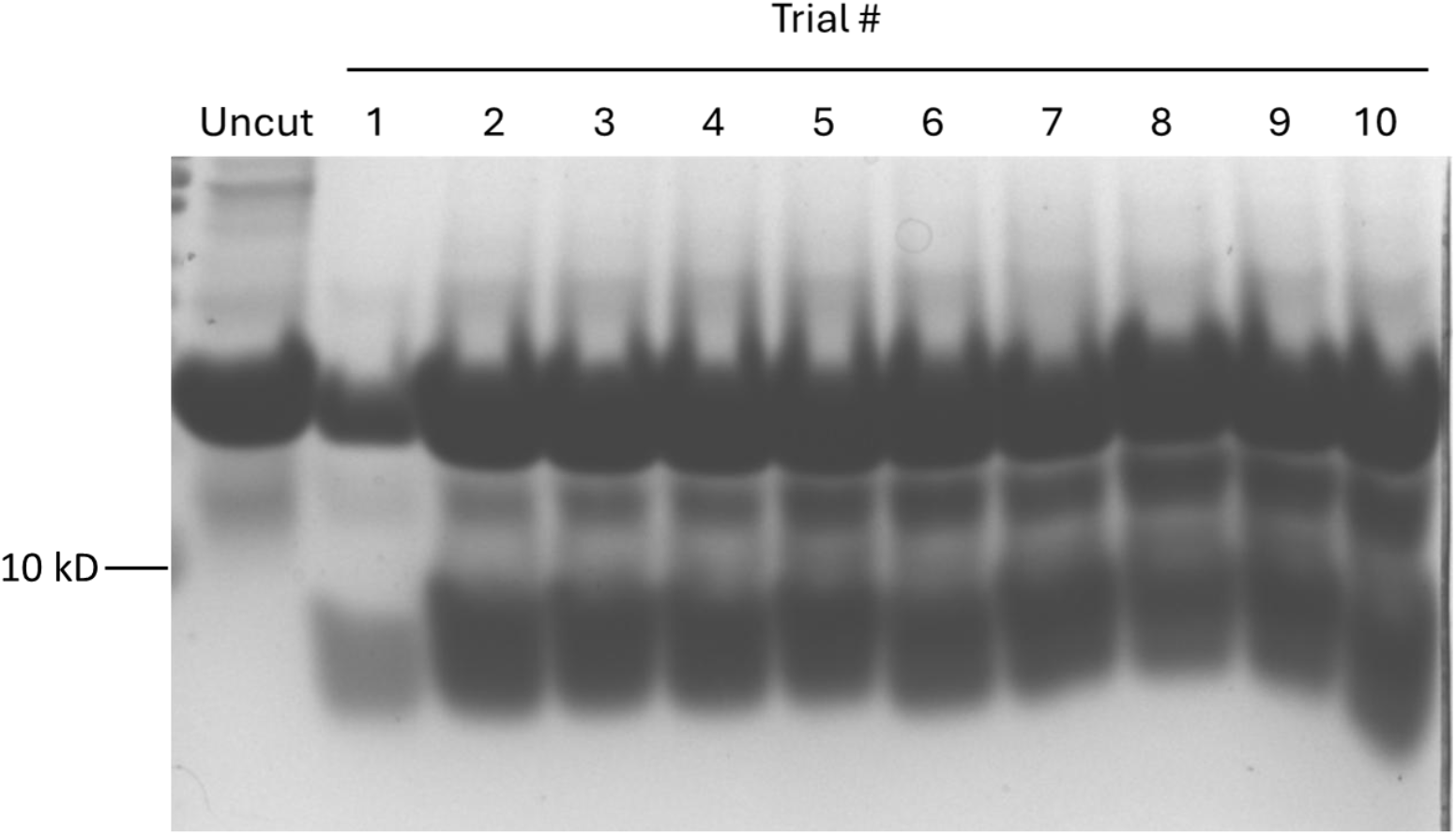
Reusability of Immobilized SUMO Protease in P113 cleavage. 18% SDS-PAGE gel demonstrating repeated use of a single aliquot of immobilized SUMO protease through ten successive applications of SUMO-P113 substrate.

We further isolated the cleaved peptide product by HPLC and confirmed its identity by MALDI-TOF mass spectrometry (**Fig. 7**). HPLC fractions absorbing at 214 nm (**Fig. 7A**) were collected and lyophilized to dryness and subsequently reconstituted in distilled deionized water. Aliquots of these fractions were analyzed on an 18% SDS-PAGE gel (**Fig. 7B**) to identify the peptide containing fraction. The peptide fraction was analyzed by MALDI-TOF mass spectrometry (**Fig. 7C**) and found to have the expected molecular weight (1564.832 daltons; calculated mass 1564.82 daltons). ESI mass spectrometry analysis of isolated peptide also contained the expected mass fragments of 783.2 (M + 2H)^2+^ and 1565.1 (M+H)^+^.

**FIGURE 7.**
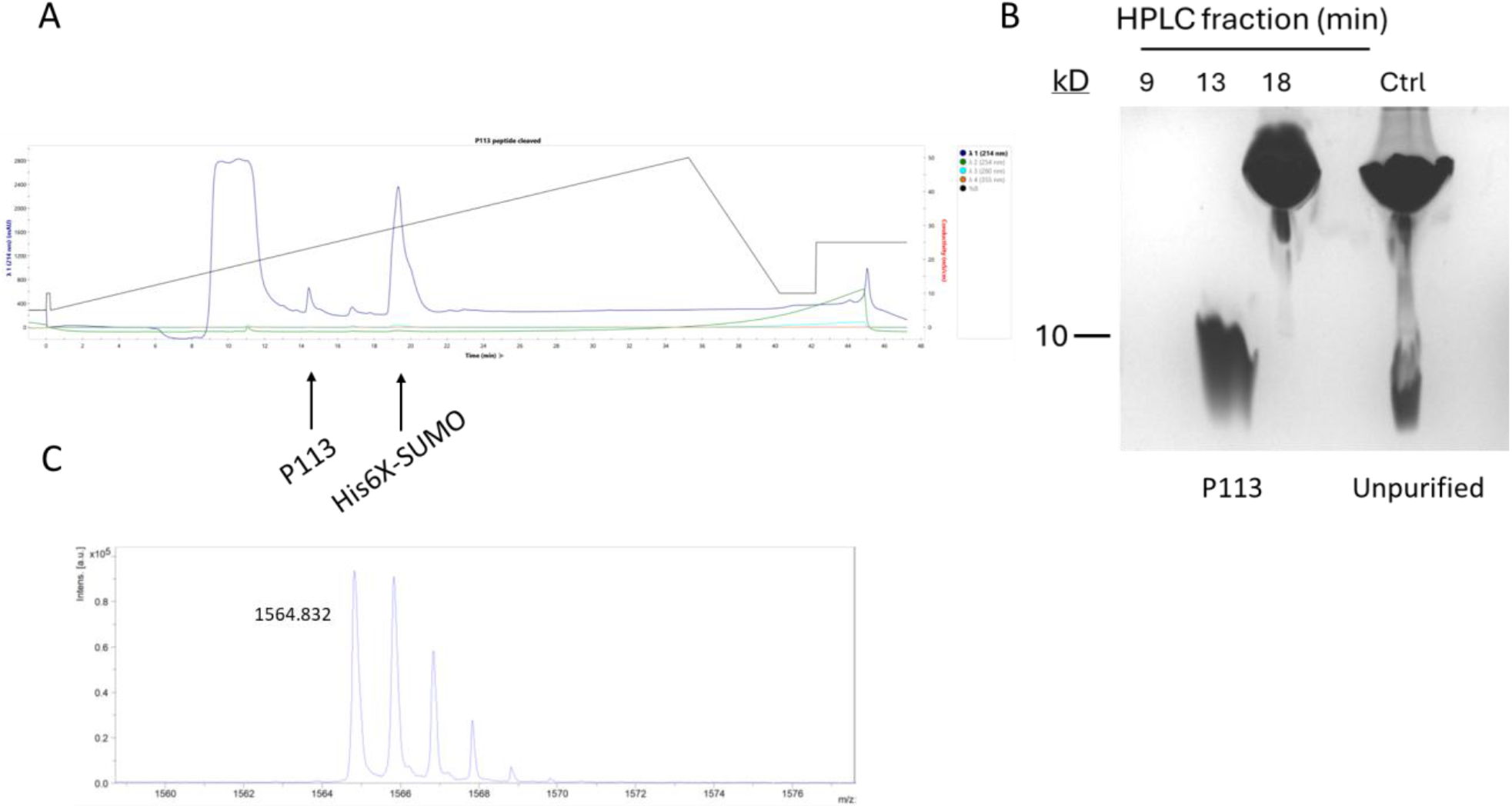
HPLC and MALDI of recombinant P113 peptide. A. Reverse phase HPLC of SUMO-P113 peptide post-overnight incubation with immobilized SUMO protease. Peaks were monitored at 214 nm. B. 18% SDS-PAGE gel of lyophilized HPLC fractions. Peak collected between 13 and 14 minutes contained the P113 peptide and peak collected between 18 and 19 minutes contained the cleaved His6X-SUMO tag fragment. C. MALDI of the peptide fragment recovered between 13 and 14 minutes has the expected intact molecular weight (1564.832 daltons; calculated mass 1564.82 daltons).

During our investigations, we observed that the production of peptide from soluble SUMO-P113 was variable, and that much of the SUMO-P113 fusion was present in the insoluble pellet fraction (**Fig. 8**). This occurred with expression in rich media (Terrific broth (TB), 2XYT, Dynamite) and to a lesser extent in LB media (**Fig. 8A**). As a result, recovery of peptide from the insoluble fraction was added to the workflow, which improved the reliability of our yields (**Fig. 8C**). We further optimized yield using a urea-based total protein extraction protocol that also streamlined our overall peptide extraction and purification protocol. As shown in **Fig. 8C**, the highest yield obtained with our system thus far was 4.7 mg/L of culture with 2XYT media and the total protein extraction method. This yield is comparable to a previously reported BrCN-based method for P113 peptide production in *E. coli* (29), which generated 4 mg/L of peptide. We note that with additional modifications to our system, including possible redesign of the SUMO-peptide fusion construct, it may be possible to further increase the yield. In addition, although our strategy produces modest yields, it provides notable “green” benefits versus SPPS by eliminating the use of DMF and peptide coupling and deprotecting reagents (30). Moreover, this strategy eliminates 90% of trifluoroacetic acid usage by restricting TFA usage to HPLC solvents only. Increasingly considered the most fundamental form of PFAS, TFA is increasingly under scrutiny for its impact on human health and is likely to fall under more regulatory control (31). The same can be said for DMF, which is already undergoing increased regulation in Europe (32). As a result, recombinant peptide product strategies may be increasingly more attractive in the future if a more restrictive regulatory environment changes the economics of SPPS.

**FIGURE 8.**
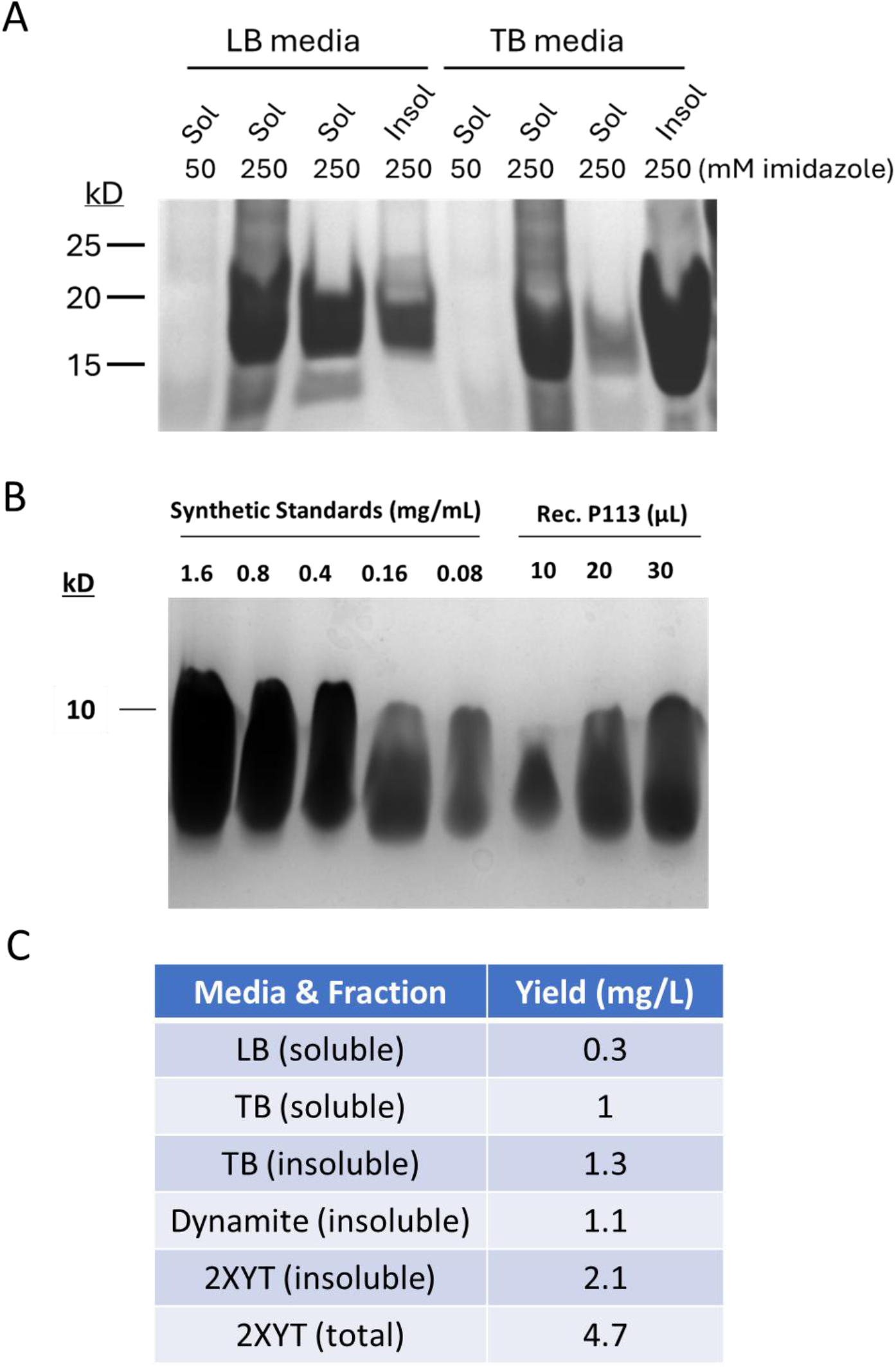
Quantification of recombinant P113 from different media types and fractions. A. SDS-PAGE gel comparing soluble and insoluble fractions recovered from expression of SUMO-P113 construct in LB and TB media. B. 18% SDS-PAGE gel comparing synthetic P113 peptide to recovered recombinant P113. C. Yields of P113 peptide obtained from soluble, insoluble, and total protein fractions in various media types. 2XYT (total) yield is from the urea-based total protein prep.

Finally, the antifungal susceptibility of recombinant P-113 was evaluated against Candida strains (**Fig. 9**). The recombinant peptide exhibited inhibitory activity against the fluconazole-resistant Candida albicans strain (ATCC 321182) at a concentration of 100 μM, and against the non-albicans Candida strain Candida tropicalis (ATCC 750) at 10 μM. The minimum fungicidal concentration (MFC) was not reached within the range of concentrations tested for either strain. Therefore, at the minimum inhibitory concentration (MIC), the recombinant peptide demonstrated a fungistatic effect against both C. albicans ATCC® 321182 and C. tropicalis ATCC® 750 (**Fig. 10A**). This is consistent with recent reports of inhibitory but non-lethal P113 activity in growth assays (33).

**FIGURE 9.**
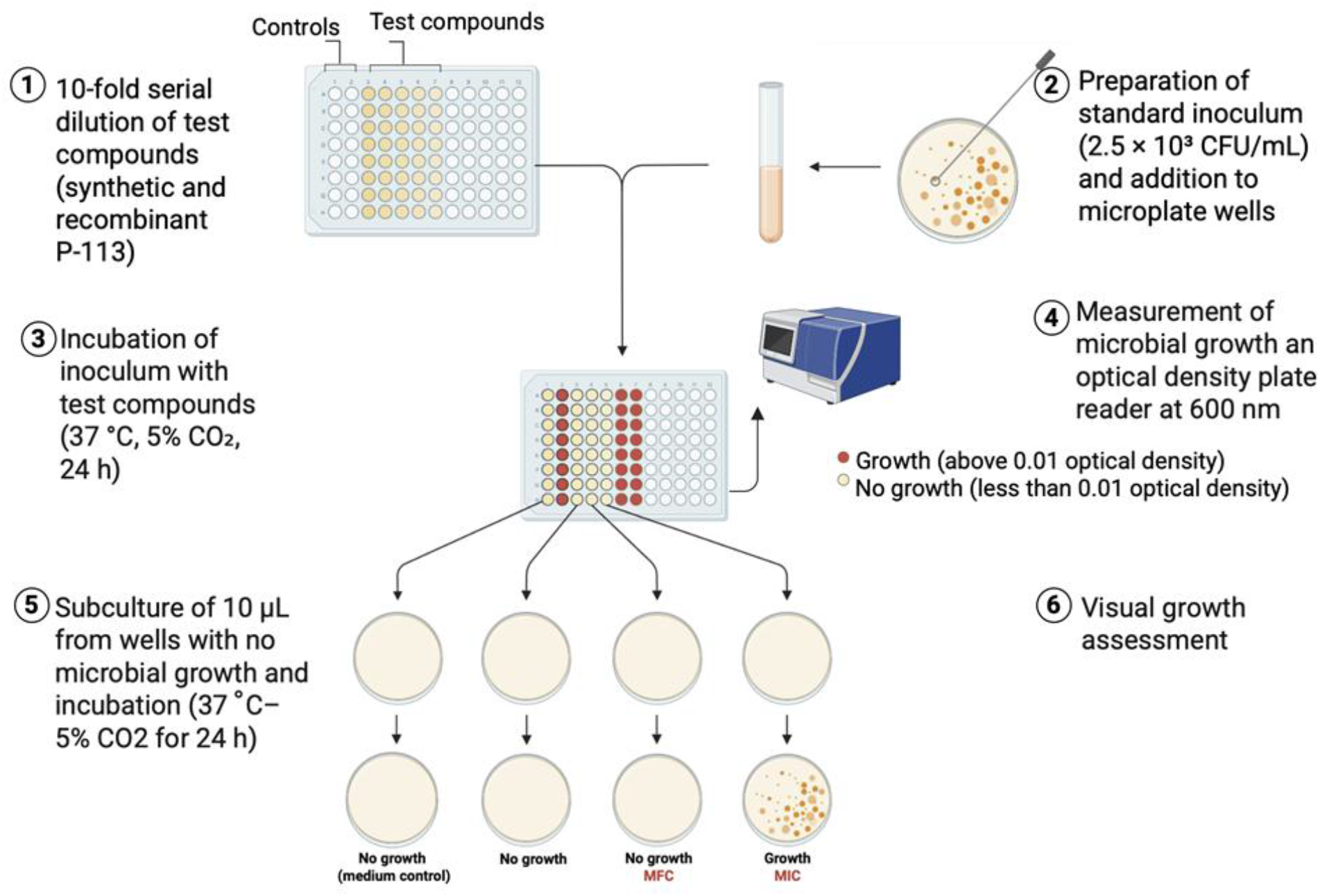
Candida spp. susceptibility test workflow. 1. Synthetic and recombinant P-113 peptides are subjected to 10-fold serial dilution in a microplate, and control wells (medium control and vehicle control) are also included. 2. A standard inoculum of Candida spp. (2.5 × 10^3^ CFU/mL) prepared in RPMI-1640 medium is added to the wells. 3. Microplates are incubated at 37 °C with 5% CO_2_ for 24 h. 4. Microbial growth is assessed by measuring optical density at 600 nm, and the lowest concentration that inhibits growth is defined as the minimum inhibitory concentration (MIC). 5. Aliquots (10 µL) from wells showing no microbial growth are subcultured onto Sabouraud Dextrose Agar plates and incubated at 37 °C with 5% CO_2_ for 24 h. 6. Microbial growth is visually evaluated to determine the minimum fungicidal concentration (MFC).

**FIGURE 10.**
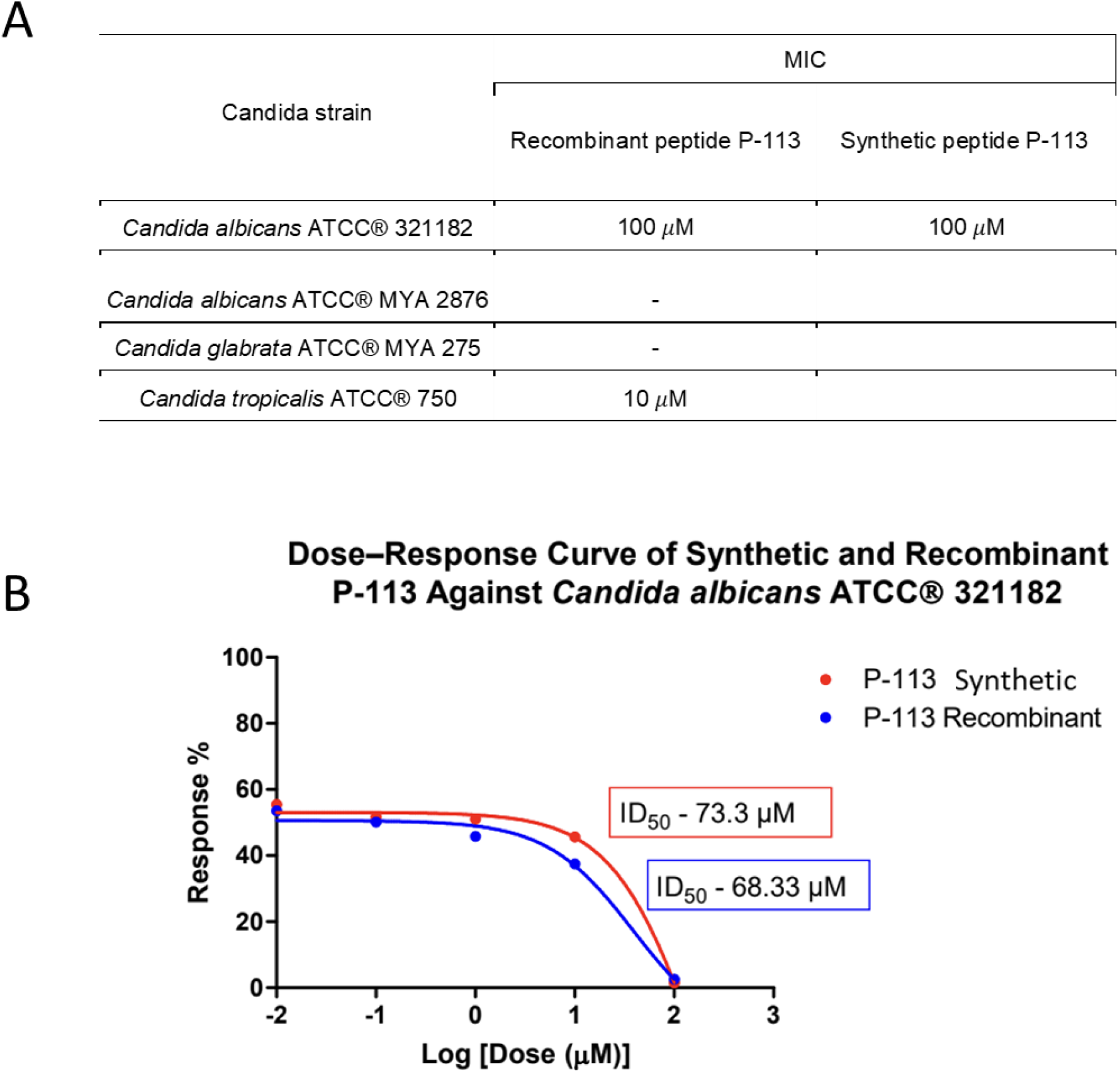
Minimum Inhibitory Concentration (MIC) and Dose–response curve of Recombinant and Synthetic peptide P-113 against fluconazole-resistant Candida albicans® ATCC 321182. A. MIC of Recombinant peptide P-113 against Candida strains (Candida albicans ATCC® 321182; Candida albicans ATCC® MYA 2876; Candida glabrata ATCC® MYA 275; Candida tropicalis ATCC® 750); MIC of Synthetic peptide P-113 against Candida albicans ATCC® 321182. Cells marked with “-” indicate microbial growth at the tested concentrations, suggesting that higher concentrations should be evaluated. B. Both peptides exhibited a concentration-dependent antifungal effect, with increasing doses leading to reduced cell viability. The toxic dose range was identified based on the significant decrease in fungal growth, highlighting comparable antifungal activity between recombinant and synthetic peptides.

The inclusion of the C. albicans ATCC® 321182 strain is particularly relevant, as it was isolated from an HIV-positive patient and displays resistance to fluconazole, a frontline antifungal agent widely used in clinical practice (34). In this context, and to enable a direct comparison, the susceptibility of this strain to the synthetic P-113 peptide was also investigated. The synthetic peptide exhibited inhibitory activity at the same concentration (100 μM), similar to the recombinant counterpart (**Fig. 10A**).Consistent with previous studies comparing synthetic and recombinant P113 peptides, no activity differences between synthetic and recombinant peptide sequences were identified (14, 16).

Furthermore, dose–response curve analysis (**Fig. 10B**) was performed to compare the antifungal activity of both peptides in terms of their ability to reduce fungal cell viability across a range of concentrations. Although both peptides exhibited a similar response profile, the recombinant peptide demonstrated greater potency, as indicated by a lower inhibitory dose (ID_50_ = 68.33 μM) compared to the synthetic peptide (ID_50_ = 78.3 μM).

## MATERIALS AND METHODS

### Construct design, expression, and purification

Immobilized SENP1 was designed based on prior work with immobilized PKAcs and TEV protease fusions (24, 25). Briefly, a HaloTag protein was connected to a SENP1 protein with a flexible GGSD repeat linker. A 6X His tag was included at the C-terminus of the fusion protein. For the SUMO tagged P113 sequence, a protein fusion with a 6X His tag, followed by a SUMO tag fused directly to the P113 sequence was created. The genes encoding the fusion protein sequences were codon optimized, synthesized, and cloned into pET28a(+) vector by Twist Bioscience (South San Francisco, CA). The plasmids were subsequently transformed into BL21-Codon Plus RIPL (Agilent Technologies) competent cells. ***Protein expression***. Freshly struck cultures of competent cells were used to inoculate overnight starter cultures (LB-Kan) supplemented with glucose (final concentration: 0.2% w/v), and grown overnight with shaking (220 rpm) at 30 °C. The following day, 20 mL of overnight culture was used to seed 1 L of cell culture media in 2L baffled flasks, then grown to OD_600_ = 0.5-0.6 (LB; OD_600_ = 1.1 – 1.2 for rich media); at this point the temperature was lowered to 17 °C and protein expression was induced with IPTG (final concentration 0.5 mM). After induction (16 - 48 h), the cells were pelleted via centrifugation (4000 RPM, 4 °C) for downstream processing. Cell pellets were lysed with 10 mL B-PER (ThermoFisher) containing 1 mM PMSF protease inhibitor. Lysates were then clarified by centrifugation (14,000 x g, 20 min, 4 °C) and the supernatant collected for purification via IMAC. The supernatant from clarified cell lysate was applied to a gravity flow column containing HisPur Cobalt Resin (Thermo Scientific), washed with PBS, and then eluted with increasing concentrations of imidazole in PBS. To recover insoluble protein fractions, the lysed cell pellet was resuspended with vortexing in 50 mL of a 1:10 dilution of B-PER reagent and then re-pelleted by centrifugation (14,000 x g, 20 min, 4 °C). This process was repeated two additional times. The washed pellet was then resuspended in 5 mL of ice cold denaturing buffer (6M GdHCl/50 mM HEPES buffer, pH 7.5) and allowed to sit on ice overnight. Additional denaturing buffer (3 mL) was then added prior to clarification of the denatured protein solution via centrifugation (14000 rpm for 30 min at 4 deg C). The clarified supernatant was then applied directly to a cobalt column to capture the solubilized protein. After column capture, the beads were washed with PBS (1 × 5 mL) to remove residual denaturing buffer and eluted with 250 mM imidazole elution buffer (2 × 7 mL fractions). These fractions were combined and dialyzed against PBS (overnight, 4 °C, Snakeskin dialysis tubing) prior to cleavage with the immobilized SENP1 enzyme. ***Enzyme immobilization***. Immobilized SENP1 fusion proteins were prepared using purified HaloTag-fusion protein and Magne-HaloTag beads (Promega). In a centrifuge tube, a 1 mL slurry of beads (5 mg) was pelleted using a magnetic separator and the suspension solution was removed. Beads were then washed with PBS (2 × 1 mL) and resuspended in 1 mL of PBS. This bead suspension is then added dropwise to a 20 mL solution of purified fusion protein (1.0 mg/mL concentration) and incubated on rocker for 2h at room temperature. The loaded beads were then pelleted by magnet, supernatant removed by pipette, and subsequently washed with PBS (3 × 5 mL). The loaded beads were resuspended to a final volume of 4 mL in a glycerol storage buffer (20 mM Tris, 300 mM NaCl, 1 mM EDTA, 50% glycerol, pH 8.0). ***Immobilized SENP1 Cleavage***. To the dialyzed samples, DTT was added final concentration of 0.5 mM (from a 1 M stock solution) followed by 50 uL of resuspended SENP1 beads in a screw top tube. Tubes were secured on their sides and allowed to rock gently overnight at room temperature. The next day, SENP1 beads were collected by magnet and the cleavage mixture transferred to a clean screw top tube. A small aliquot (200 uL) was taken for analysis by SDS-PAGE (18% gel).

The remaining solution was frozen and lyophilized. The recovered beads were washed with PBS and placed in glycerol storage buffer for later reuse. ***Protein separation***. IMAC fractions were resolved on 10% SDS-PAGE gels in MOPS buffer and stained with Simplyblue SafeStain (Invitrogen); Enzymatic cleavage reactions were resolved on 18% SDS-PAGE gels in MOPS buffer. ***Total Protein Extraction with Urea***. Freshly struck cultures of competent cells were used to inoculate overnight starter cultures (LB-Kan) supplemented with glucose (final concentration: 0.2% w/v), and grown overnight with shaking (220 rpm) at 30 °C. The following day, 20 mL of overnight culture was used to seed 1 L of 2xyt cell culture media in 2L baffled flasks, then grown to OD_600_ = 1.1 – 1.2 at 37 °C; at this point the temperature was lowered to 30 °C and protein expression was induced with IPTG (final concentration 0.5 mM). After induction (18 h), the cells were pelleted via centrifugation (4000 RPM, 4 °C) for downstream processing. Cell pellets were lysed with 10 mL B-PER complete (ThermoFisher) supplemented with 1 mM PMSF protease inhibitor and 6.4 M urea. Lysates were then clarified by centrifugation (14,000 x g, 20 min, 4 °C) and the supernatant collected for purification via IMAC. The supernatant from clarified cell lysate was applied to a gravity flow column containing HisPur Cobalt Resin (Thermo Scientific), washed with PBS (10 mL), and then eluted with 250 mM of imidazole in PBS in three fractions (2 × 10 mL and 1 × 5 mL). These fractions were combined and diluted 1:1 with PBS. DTT was added to the solution to a final concentration of 0.5 mM, and 200 μL of resuspended immobilized SENP1 enzyme beads added. The solution was allowed to incubate overnight at room temperature with gentle rocking. The next day, the cleaved mixture was removed from the immobilized enzyme beads, frozen, and lyophilized prior to further purification with RP-HPLC.

### HPLC and Mass spectrometry

HPLC separations were conducted using a Bio-Rad NGC 10 Plus chromatography system with an Agilent C-18 Prep column and detection at 214 nm. Peptides were separated with gradient elution running from 10% ACN/0.1%TFA to 100 % ACN/0.1% TFA over 35 minutes with a flow rate of 5.0 mL/min. After HPLC separation, fractions were frozen and lyophilized for later analysis. Electrospray mass spectra were acquired on an API 3200 triple quad mass spectrometer and MALDI mass spectra were acquired on a Bruker UltrafleXtreme MALDI-TOF mass spectrometer.

### Candida spp. susceptibility test methodology

*Microorganisms*. The following standard ATCC (American Type Culture Collection®) reference yeast of Candida were used: C. albicans ATCC® 321182, C. albicans ATCC® MYA 2786, C. glabrata ATCC® MYA 275 and C. tropicalis ATCC® 750. *Determination of Minimal Inhibitory Concentration (MIC) and Minimal Fungicidal Concentration (MFC)*. The microdilution method (35, 36) was used to determine the MIC and MFC of the Candida strains according to CLSI guidelines (M27-A2) (Clinical and Laboratory Standards Institute (CLSI). Roswell Park Memorial Institute Medium—RPMI-1640 (Corning®1) was inserted into the wells, followed by different concentrations of Synthetic Peptide P113 (100 to 0.01μM) and Recombinant Peptide P113 (100 to 0.01μM). Lastly, fungal suspension (2.5 × 103 colony forming units - CFU/ml) were added to the wells. Wells containing DMSO 1%, inoculum and medium were used as the vehicle control. Wells containing only culture medium were used as the medium control.

Plates were incubated at 37 °C– 5% CO2 for 24 h, and microbial growth was measured using an optical density plate reader. Wells showing no visible growth (less than 0.01 optical density units at 600nm) were considered inhibited. Among these wells, the lowest concentration capable of inhibiting microbial growth was defined as the minimum inhibitory concentration (MIC). Subsequently, 10 μL aliquots from wells corresponding to concentrations equal to/higher than the MIC were subcultured onto Sabouraud Dextrose Agar (Kasvi®) and incubated at 37 °C with 5% CO_2_ for 24 h. After incubation, visual growth was analyzed to determine the minimum fungicidal concentration (MFC).

Additionally, based on the optical density data obtained from the antifungal susceptibility assay, dose–response curves were generated for both recombinant and synthetic P-113 peptides. Fungal growth inhibition was expressed as a function of peptide concentration, and the data were fitted using nonlinear regression analysis to determine the inhibitory dose (ID_50_) values for each compound.

## CONCLUSIONS

In this work, we have demonstrated the application of an immobilized enzyme in a recombinant peptide production method. The immobilized enzyme can be used for repeated cleavage reactions (at least 10X) and retains activity long after its initial preparation (> 6 months) in cold storage. Importantly, the immobilized enzyme is a viable substitute for cyanogen bromide as it 1) produces the native enzyme sequence and 2) can generate quantifiable yields of P113 that enable molecular characterization and bioassay. While the yields are modest compared to what is readily achieved with solid phase peptide synthesis, the reduced environmental impacts of this approach are compelling. We are currently working on additional optimizations to this method to increase peptide yield. In the future, we plan to use this approach in the production of additional P113 variants and in the production of other non-Histatin peptide sequences.

## ACKNOWLEDGEMENTS

We acknowledge Mr. Thomas Cope (laboratory of Prof. Marcey Waters, University of North Carolina – Chapel Hill) for providing the synthetic P113 peptide standard. We also acknowledge Dr. Ninad Doctor (ECU Department of Chemistry) for assistance with electrospray mass spectrometry and Dr. Brandi Ehrman (University of North Carolina – Chapel Hill) for performing the MALDI mass spectrometry.

